# Remnants of the Balbiani body are required for formation of RNA transport granules in *Xenopus* oocytes

**DOI:** 10.1101/2020.05.27.119487

**Authors:** Chao Yang, Gena M. Wilson, Matthew M. Champion, Paul W. Huber

## Abstract

Fragmentation of the Balbiani body in *Xenopus* oocytes engenders a region extending from the germinal vesicle (GV) towards the vegetal pole that is enriched in mitochondria. This area is later transversed by RNA that is being localized to the vegetal cortex. Inhibition of mitochondrial ATP synthesis prevents the perinuclear formation of these RNA transport granules that can be reversed by the nonhydrolyzable ATP analog, adenosine 5’-(βγ-imido) triphosphate. The protein composition and sensitivity of the transport granules to hexanediol indicate that they are liquid phase condensates. Mitochondria in the remnants of the Balbiani body produce a region of elevated ATP that appears to act as a hydrotrope to support the perinuclear phase transition leading to granule formation.

**Summary:** Mitochondria in the remnants of the Balbiani body produce elevated levels of ATP required for the formation of liquid phase condensates containing RNA transport granules.

---

The Balbiani body (Bb), an organelle comprised of mitochondria, endoplasmic reticulum, and RNA, is found in the oocytes of most organisms (*1*). In *Xenopus*, the structure is initially positioned adjacent to the oocyte nucleus, also known as the germinal vesicle (GV), facing the vegetal pole. The Bb eventually extends towards the vegetal cortex and delivers germ plasm determinants including *nanos1* and *wnt11* mRNAs, after which it fragments, leaving a cytoplasmic region rich in mitochondria (*2*). Shortly after the breakdown of the Bb, a second group of mRNAs begins to move to the vegetal cortex through a process dependent on the cell cytoskeleton (*3, 4*). This latter gradient of localized mRNAs, which include *Vg1* and *VegT*, ultimately contributes to axis formation in the embryo.

Oocytes from transgenic frogs that express green fluorescent protein (GFP)-tagged mitochondrial outer membrane protein 25 (*5*) were used to examine more closely the relation between the remnants of the dispersed Bb and RNA transported through the late pathway to the vegetal hemisphere. A 453-nucleotide RNA containing the vegetal localization element (VLE) of *Vg1* mRNA is efficiently transported to the vegetal cortex when injected into stage II/III oocytes and is used to track movement of RNA through this pathway (*6, 7*). The region formerly occupied by the Bb retains a high concentration of mitochondria that exhibit a striking colocalization with the *Vg1* RNA transport particles, especially the perinuclear “cup” structure where the RNA first enters the cytoplasm (Fig. 1A).

**Fig. 1.**
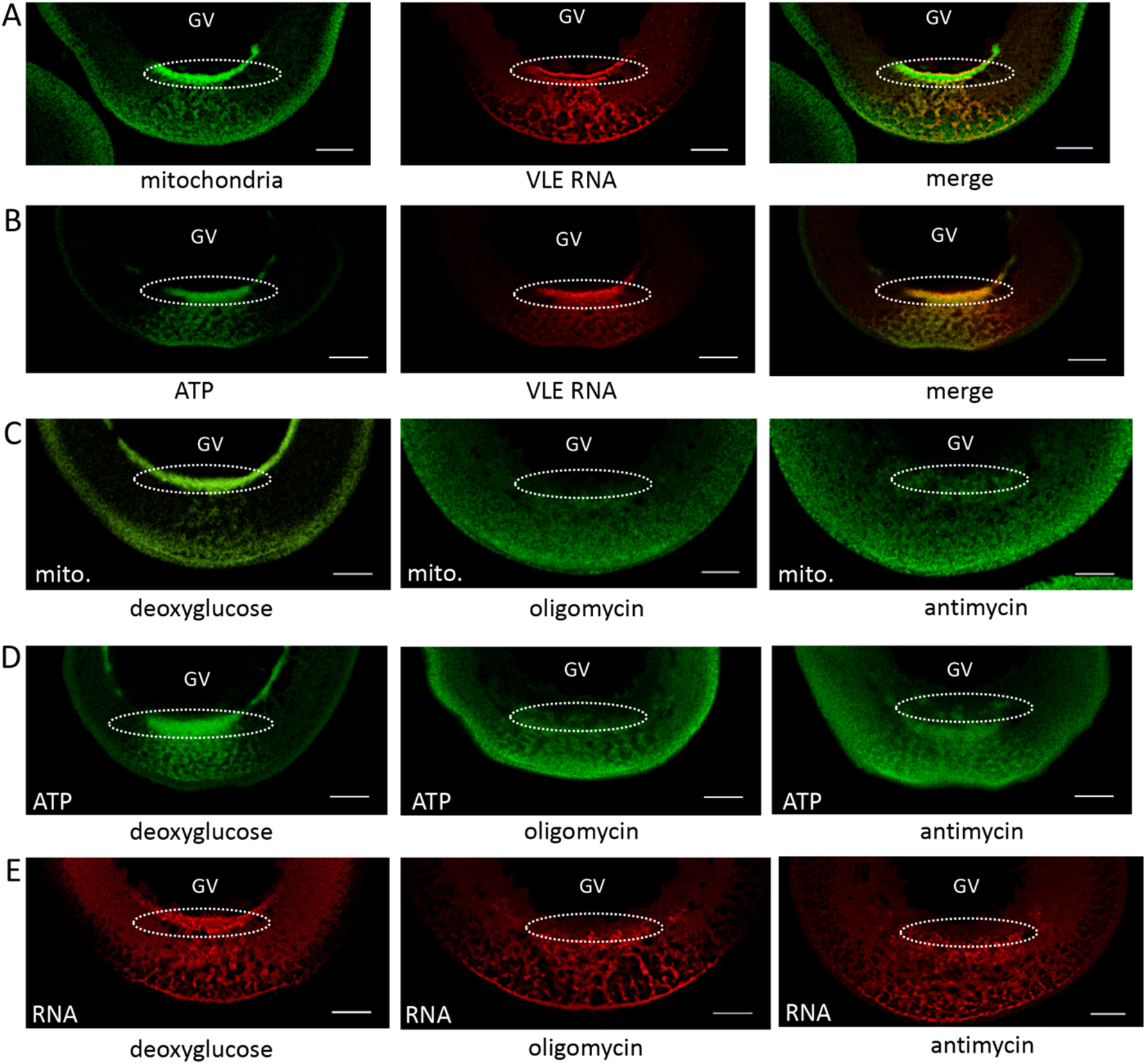
Remnants of the Balbiani body generate a region of elevated ATP that supports formation of RNA transport granules. **(A)** The area formerly occupied by the Bb is enriched in mitochondria (*left*); VLE RNA being transported to the vegetal cortex (*center*) traverses the same area (*right*). The perinuclear “cup” structures where the RNA exits the vegetal side of the GV are enclosed in circles. **(B)** The clustered mitochondria of the Bb generate a region of elevated ATP concentration that is coincident with VLE RNA. **(C-E).** Inhibition of glycolysis (*left*) has no effect on the clustering of mitochondria **(C)**, localized ATP production **(D)**, or RNA transport **(E).** Inhibition of mitochondrial ATP synthesis with oligomycin (*center*) or antimycin (*right*) eliminates the clustered region of mitochondria **(C)** and ATP (**D)**, and suppresses formation of the RNA granules at the perinuclear cup **(E)**. Scale bars = 50 μm.

There are recent reports of subcellular enrichment of mitochondria in regions where there is high demand for ATP that, in one case, supports local protein synthesis during dendritic synaptic plasticity (*8*) or actin network growth in another case (*9*). It appeared possible that the mitochondria enriched in the vegetal hemisphere might be capable of generating locally elevated amounts of ATP. Therefore, we employed the biosensor protein, ATeam (*10*), to visualize ATP levels in oocytes and detected regions of increased ATP within the area containing the remnants of the Bb, especially the cup structure in the vegetal perinuclear region (Fig. 1B). In order to relate the enriched region of ATP to mitochondrial activity we treated oocytes with the inhibitors antimycin A (inhibitor of complex III of the electron transport chain), oligomycin (inhibitor of ATP synthase) or deoxyglucose (inhibitor of glycolysis). Whereas antimycin and oligomycin eliminated the areas of elevated ATP, deoxyglucose had no detectable effect, implicating mitochondria from the disseminated Bb fragments in generating locally high concentrations of the nucleotide (Fig. 1C and D).

We next addressed the question whether the elevated amounts of ATP are needed for RNA localization activity. Notably, the greatest effect of antimycin A and oligomycin is on the formation of the perinuclear cup structure with a lesser effect on movement of localization granules approaching the vegetal cortex (Fig. 1E). Deoxyglucose had no effect on any aspect of *Vg1* RNA localization. These results indicate that basal levels of ATP are generally sufficient to support motor-dependent movement of RNP particles along the cytoskeleton, but that the formation of the cup structure at the nuclear periphery, in particular, requires the elevated amounts of ATP generated by the mitochondria of the Bb fragments. It appears that the increased amount of ATP in the region formerly occupied by the Bb helps maintain the structure of the RNP particles moving to the vegetal cortex, since the number of particles was often diminished in oocytes treated with antimycin A or oligomycin.

There is evidence that *Vg1* RNP complexes undergo some type of remodeling event upon entering the cytoplasm that affects protein-RNA and protein-protein interactions (*11*); however, it is not known whether this process requires ATP. A number of ATP-dependent RNA helicases mediate changes in RNP structure before and after nuclear export to the cytoplasm (*12*). Thus, it appeared possible that the elevated levels of ATP are needed to support RNA helicase activity that might be required for the formation of active transport complexes at the perinuclear cup. Alternatively, ATP, at millimolar concentrations, can act as a hydrotrope that stabilizes protein structure (*13, 14*), and in this instance could be maintaining the solubility of proteins in the region of the cup structure, thus, enabling formation of the transport particles.

To assess the requirement for ATP hydrolysis, we tested the ability of a nonhydrolyzable analog of ATP, adenosine 5’-(βγ-imido) triphosphate (AMP-PNP), to rescue formation of the cup structure in the presence of oligomycin or antimycin A (Fig. 2). AMP-PNP was co-injected with VLE RNA to yield a final *in vivo* concentration of approximately 4 mM and the oocytes cultured for 12 hr. The nonhydrolyzable analog was able to maintain cup formation and stabilize *Vg1* particles in oocytes incubated with either oligomycin or antimycin A. ATP concentration is not uniform in amphibian oocytes, being lowest in the vegetal cytoplasm (1.9 mM) which is just below the effective concentration range (2 - 8 mM) in which ATP acts to prevent protein aggregation (*13, 15*). In contrast, the concentration of ATP in the *Xenopus* oocyte GV is 6 mM. Rescue of cup formation by AMP-PNP indicates that the high concentration of ATP generated by mitochondria clustered at the nuclear periphery is necessary to maintain protein solubility as the RNP complexes enter the cytoplasm. Indeed, the hydrotropic action of ATP has been shown to solubilize protein aggregates in *Xenopus* oocyte nucleoli (*16*). The need for elevated amounts of ATP to support helicase activity cannot be fully discounted; however, the sub-millimolar Kd values for ATP binding to DEAD-box helicases makes this situation unlikely (*17*). Rescue by the nonhydrolyzable analog also eliminates protein phosphorylation as the basis of the inhibition by oligomycin and antimycin A (*18*).

**Fig. 2.**
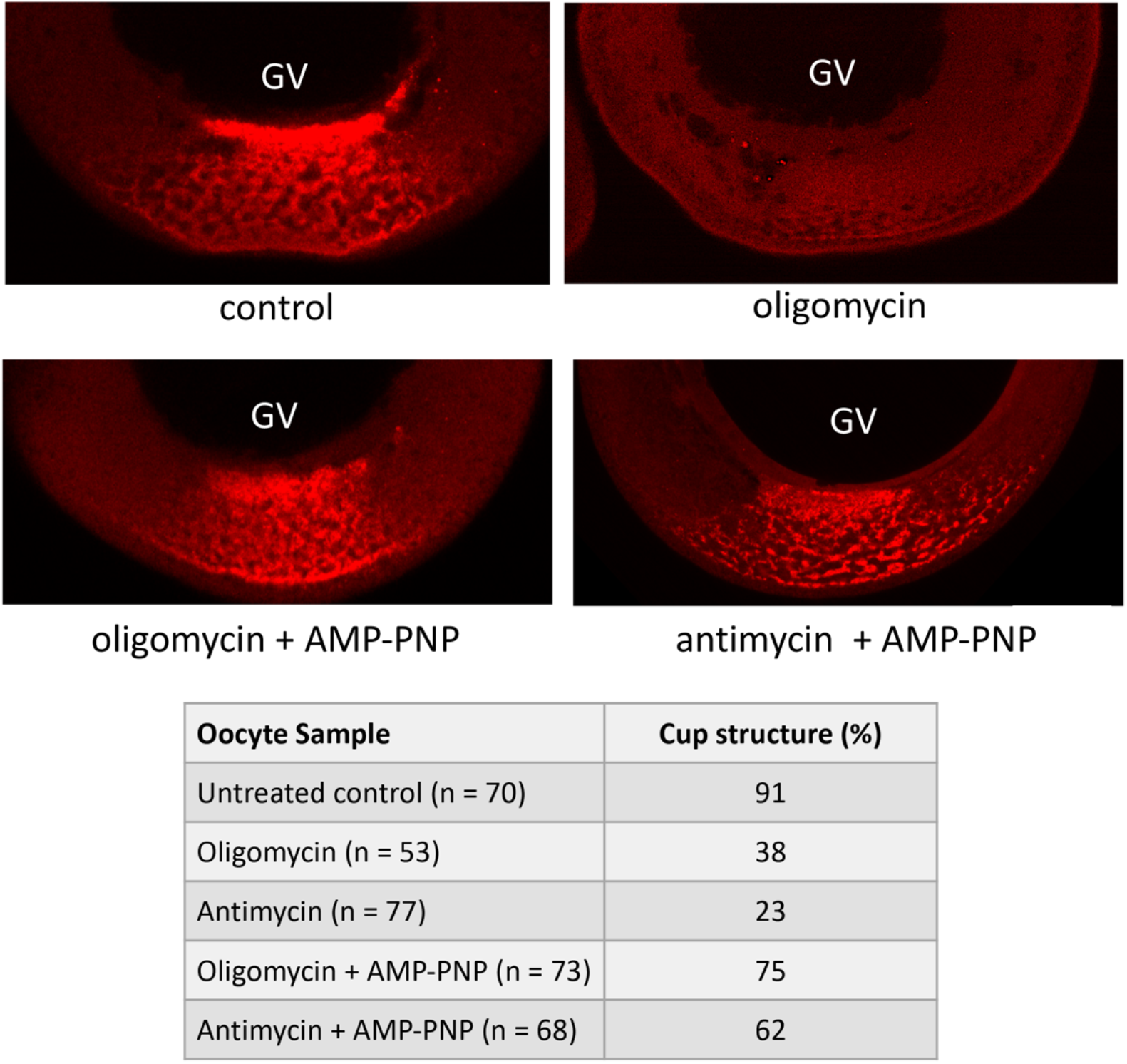
Rescue of RNA granule formation with a nonhydrolyzable analog of ATP. Oocytes were injected with VLE RNA and placed in medium containing either oligomycin or antimycin. Coinjection with adenosine 5’-(βγ-imido) triphosphate (AMP-PNP) to give a calculated intracellular concentration of 4 mM is sufficient to restore the formation of the perinuclear cup structure and distinct RNA granules.

The cup structure appears to be a region where localized RNAs such as *Vg1* are assembling into larger structures prior to movement towards the vegetal cortex. RNA transport is known to occur in phase-separated liquid droplets in several cell types (*19*). These structures, similar to stress granules and P-bodies, are sensitive to dispersion in the presence of 1,6-hexanediol. Oocytes injected with VLE RNA were cultured for 16 hours and then incubated in 5% 1,6-hexanediol for 1 hour. Much of the pattern of VLE RNA is lost upon this treatment, but the effect is especially noticeable at the perinuclear cup structure, which is entirely eliminated (Fig. 3A). Coalescence of the VLE transport particles and marked enlargement of the cup upon inhibition of transport by microtubule depolymerization with nocodazole provides additional evidence that these structures are a phase-separated condensate (Fig. 3B and Fig. S1). Most injected VLE RNA typically clears the perinuclear region after 24 hr, but in the presence of nocodazole, a considerable amount of the RNA remains in an expanded cup structure even at 48 hr. This behavior is notably similar to the effect of nocodazole on RNA granules in zebrafish germ cells (*20*).

**Fig. 3.**
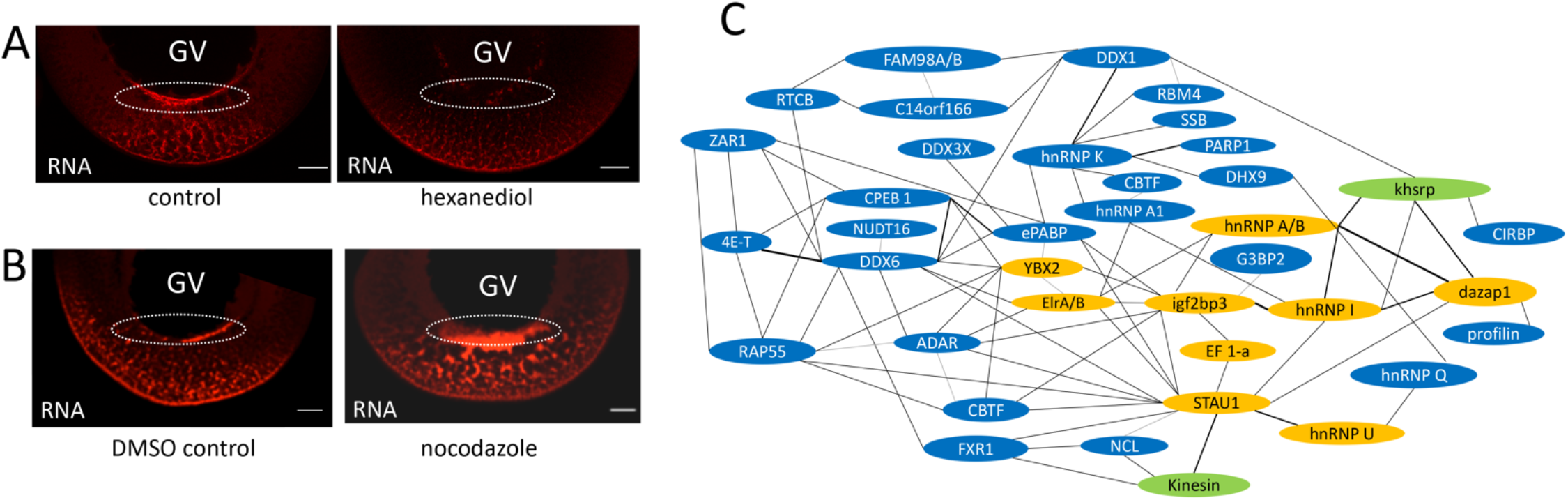
The VLE RNA transport granules are a phase-separated biocondensate. **(A)** Oocytes were taken 16 hr after injection with VLE RNA and treated with 5% 1,6-hexanediol for 60 minutes and then fixed. **(B)** Following injection with VLE RNA, oocytes were placed in culture medium containing DMSO (vehicle control) or nocodazole for 48 hr and then fixed. **(C)** Protein-protein interaction map constructed from proteins identified in the VLE RNP complex affinity purified from oocyte extract. Heavy lines indicate direct interactions and light lines connect proteins for which a direct interaction has not been established. Gold background indicates proteins previously identified as binding directly to VLE RNA; blue background indicates newly identified proteins; VgRBP71 (khsrp), which binds directly to Vg1 RNA, and kinesin were not identified in the MS samples. Scale bars = 50 μm.

Several proteins that bind to the localization element of Vg1 mRNA have been identified (*4, 6, 21–29*) and most have also been identified in macromolecular RNP granules that undergo phase separation (*30–36*). In order to obtain a more comprehensive inventory of the proteins that comprise the *Vg1* RNP particle and to compare its composition to other phase-separated RNP granules, we affinity purified complexes that form on biotin-tagged VLE RNA in oocyte extract. Proteins in the affinity purified complexes were identified by tandem mass spectrometry (Fig. 3C, Table S1).

In addition to the known VLE-binding proteins, most of the newly identified proteins are those commonly found in non-membrane RNP structures (*30–34, 36, 37*). However, a few proteins, *e.g.* zygote arrest 1, appear specific to this particle, perhaps reflecting its particular function in the oocyte. While the translation regulatory protein 4E-T and its binding partner in P-bodies, LSM14 (RAP55) (*38*) are found in the VLE complex, another major P-body component, the Ccr4-Not complex is not. Indeed, proteins actively involved in RNA degradation or regulation by miRNA are not present in the VLE complex. It is also noteworthy that the major structural protein of the Bb, Xvelo (*18*), is not found in the VLE particle. A bioinformatics analysis revealed that 39 out of the 70 most abundant proteins contact one or more other proteins in the complex (Fig. 3C). Many of the remaining proteins are components of the cytoskeleton, likely reflecting the transport activity of the complex. The composition of the VLE RNP complex is notably similar to that of RNA transport granules in neurons that also undergo liquid-liquid phase separation (*35, 39*).

Structures analogous to the Bb are universally found in animal oocytes and their formation is an early event in establishing cell polarization; however, the function of the organelle is not clear. There are phylogenetic differences with regard to whether the organelle participates in germ cell specification (*1*). In some species (frog, fish, insect), the Bb contains germ plasm, the maternal determinants that induce germ cell fate; whereas, in others (mammals, salamander), specification is inductive and independent of this organelle. Another proposed role for the Bb is to select and protect healthy mitochondria that will be passed onto progeny often after a long period of cell dormancy (*1, 18*); although, there are no details how this process might occur. Our experiments reveal another role for this organelle, at least in amphibian oocytes. The animal-vegetal polarity established by the formation of the Bb can determine the direction of RNA movement by creating an environment that sustains formation of a biomolecular condensate. This is accomplished by the mitochondria-rich remnants of the Bb that generate elevated levels of ATP, likely acting as a hydrotrope, that promote the phase transition needed for formation of the RNA transport granule. This is possibly the first example where a region of clustered mitochondria is used not to meet extraordinary energy demands (*8, 9*), but rather to initiate cellular compartmentalization.

## Acknowledgments

We thank Dr. Michelle Joyce, Mr. William Phillips, and Mr. William Archer for their assistance with the experiments; Drs. Daniel Weeks and Kevin Vaughn for expert advice; Dr. Jessica Brown for comments on the manuscript.

## Author contributions

C.Y. and P.W.H. conceived the experiments. C.Y. performed the experimental work, except the MS/MS measurements (M.M.C.), and AMP-PNP rescue experiment (G.W.). C.Y. and P.W.H. wrote the manuscript. All authors reviewed the data and commented on the manuscript.

## Supplementary Materials

Materials and Methods

Fig. S1

Table S1

References (*39–47*)

