## Supplemental Material for "Remnants of the Balbiani body are required for formation of RNA transport granules in *Xenopus* oocytes"

Materials and Methods

**Reagents**

All specialty biochemicals were purchased from Sigma-Aldrich.

**Animals**

Wild type *Xenopus laevis* were purchased from Nasco (Fort Atkinson, WI); transgenic frogs (NXR_0059) expressing eGFP fused to OMP25-mitochondria transmembrane segment (*39*) were purchased from the National *Xenopus* Resource, Marine Biological Laboratory (Woods Hole, MA). All animal experiments were performed according to protocols approved by the University of Notre Dame Institutional Animal Care and Use Committee (Protocol Number 18-02-4408)

**Plasmids**

The plasmid pBSVg1-B was linearized with *Bam*HI for synthesis of the 453 VLE RNA (*40*). ATeam1.03-nD/nA/pcDNA3 plasmid, a gift from Takeharu Nagai (Addgene plasmid #51958) was linearized with *Not*I for synthesis of ATeam mRNA (*41*).

**RNA Synthesis and Purification**

RNA was synthesized by run-off transcription in a 50 µl reaction containing rNTP (0.5 mM), 2 mM m^7^G(5’)ppp(5’)G cap analog (Promega), DTT (10 mM), linearized plasmid (1 µg/10µl), RNasin (1 U/µl,) T7 or SP6 RNA polymerase (1 U/µl, Promega) and transcription buffer (Promega). An additional 0.5 U/µl of RNA polymerase was added after 2 hr incubation for a total reaction time of 5 hr at 37 ℃. Following RNA synthesis, the DNA template was degraded by addition of RQ1 RNase free DNase (0.5 U/µl, Promega) and the mixture was incubated for 25 min at 37 °C. The reaction was stopped by adding 1 µl of 0.5 M EDTA. The volume of the RNA solution was brought up to 200 µl by adding DEPC-treated H_2_O and extracted once with an equal volume of phenol (pH 4.5) and once with equal volume of chloroform:isoamyl alcohol (24:1) at room temperature, followed with ethanol precipitation (2.5 volume of ice cold ethanol, 0.1 volume of 3 M sodium acetate) for 1 hr at -80 °C. The precipitate was washed once with 70% ice-cold ethanol and the pellet dissolved in 100 μl renaturation buffer (50 mM HEPES pH 7.4, 0.3 M KCl, 10 mM MgCl_2_), incubated for 10 minutes at 55 °C followed by slowly cooling to room temperature to allow RNA renaturation. Spin columns (1 ml bed volume) prepared with G-50 Sephadex were used to remove unincorporated nucleotides. Synthesis reactions were modified for the following RNAs: (1) RNA labeled at the 5'-terminus with biotin contained 3 mM biotin-GMP in place of m^7^G(5’)ppp(5’)G; (2) fluorophore-labeled RNA contained 0.05 mM Alexa Fluor 546-14-UTP or 488-14-UTP and 0.45 mM UTP. RNA was quantified by measuring absorbance at 260 nm using a NanoDrop 2000C.

**Oocyte Preparation, Microinjection, and Culturing**

*Xenopus* oocytes were obtained by surgically removing part of the ovary tissue and treated with 0.1 mg/ml Liberase TH collagenase (Sigma) in 25 ml OR^–^ buffer with gentle rotation at room temperature until oocytes were visibly separated from ovary matrix. Dispersed oocytes were washed 3 times with OR2^–^ buffer and 3 times with OR2^+^ buffer (OR2^-^ buffer supplemented with 1 mM MgCl_2_ and 1 mM CaCl_2_), 10 min per wash. The oocytes were manually separated according to developmental stage (*42*) and kept in OR2^+^ buffer from 3 hr to overnight to allow oocytes to recover from collagenase treatment.

A Narishige micromanipulator was used for microinjection (volumes ranged from 2 to 12 nl) of stage II/III oocytes in OR2^+^ buffer. After injection, oocytes were placed in oocyte culture medium (OCM) [50% Leibovitz’s L-15 medium, 15 mM HEPES pH 7.6, 1 mg/ml L-glutamine, 5 mg/ml vitellogenin, 1µg/ml insulin, 100 U/ml penicillin G, 100 µg/ml streptomycin, 100 µg/ml gentamicin, 50 U/ml nystatin]. OCM was made fresh and sterilized by filtration (Whatman 0.2 µm). Oocytes were cultured using 24-well plates with up to 50 oocytes per well (500 μl OCM). Culture medium was changed daily. Damaged oocytes were removed and the healthy oocytes were placed in a new well each day. The plate was placed in a hydrated container and kept at 18 ℃.

For RNA localization experiments, 150 pg VLE RNA (4 nl) Alexa Fluor 546-14-UTP labeled VLE RNA was microinjected into early stage III oocytes and the oocytes were kept in 18 ℃ for 4 to 36 hrs. For expression of ATeam protein, 250 pg mRNA (2 nl) was microinjected and oocytes cultured in OCM for 5 hr before subsequent injection of Alexa Fluor labeled VLE RNA.

**Inhibitor Studies**

For ATP inhibition experiments, oocytes were injected with VLE RNA and placed in culture medium containing either oligomycin (2 μg/ml), antimycin (1.5 μg/ml), or 10 mM 2‑deoxyglucose and 5 mM pyruvate. In rescue experiments, adenosine 5'-(β,γ-imido) triphosphate (100 mM neutralized stock) was coinjected with VLE RNA. Oocytes were fixed 12 to 16 hr after injection. For inhibition of microtubule polymerization, oocytes were injected with VLE RNA and cultured in medium containing 2 μg/ml nocodazole for 48 hr.

To confirm the depolymerization activity of nocodazole on microtubules, control or treated oocytes were fixed in MEMFA (0.1 M MOPES, pH 7.4, 2 mM EGTA, 1 mM MgSO_4_, 3.7% formaldehyde) for 1 hr and washed twice in PBST (137 mM NaCl, 2.7 mM KCl, 8 mM Na_2_HPO_4_, and 2 mM KH_2_PO_4_, pH 7.4, 0.1% Tween-20) for 10 min. Oocytes were bisected into two pieces and incubated in PBST containing 2 mg/ml BSA and 10 ug/ml β-tubulin antibody (NB600-936; Novus) overnight at 4 °C. The oocytes were washed 3 times in PBST, 3 hours for each wash at 4 °C and then incubated with Alex Fluor 546-conjugated anti-rabbit IgG (Invitrogen, 1:300 dilution in PBST with 2 mg/ml BSA) overnight at 4 °C. The oocytes were washed 3 times in PBST again for 3 hours for each wash at 4 °C. The samples were dehydrated with methanol dehydration prior to imaging.

**Microscopy**

Oocytes were fixed for 1 hr in MEMFA, washed once with OR2^+^ buffer and dehydrated by successive 5 min washes in 75% methanol-25% OR2^+^, 50% methanol-50% OR2^+^, 25% methanol-75% OR2^+^, and stored in 100% methanol at -20 ℃. Before imaging, oocytes were cleared by replacing methanol with clearing medium (2:1 benzyl benzoate:benzyl alcohol).

Images were collected on a Nikon A1R confocal microscopy with the following settings: pinhole range from 1.2-1.6, frames/second set at 1/32; scanning size at 512, integration set at “Normal”, laser power at around 100, offset at 0 and gain set at 2.69 to 2.96. To obtain the fluorescence spectra of Alexa Fluor 546 labeled RNA, oocytes were excited with 561 nm laser light, and emission from 570 to 620 nm was scanned. To capture the fluorescence spectra of the FRET-based ATP indicator with ATP bound, CFP was excited with 457 nm laser light, and emission from 500-550 nm was collected.

**Affinity Purification of the VLE RNP complex**

Oocyte extract was made by homogenizing 500 oocytes (late stage II/stage II) in 1 ml TGKED buffer (50 mM Tris pH 7.5, 25% glycerol, 50 mM KCl, 0.1mM EDTA, 0.5 mM DTT) supplemented with 1mM PMSF and 1X protease inhibitor (Roche cOmplete) with a pestle in a microcentrifuge tube. The homogenate was cleared by centrifuging at 13200 rpm for 3 min at 4 ℃. Extract was then pre-cleared with 400 μl agarose 4B beads for 1 hr at 4 ℃. Streptavidin-agarose beads (100 μl per sample, Sigma) was pre-blocked with 50 μg acetylated BSA, 30 μg yeast tRNA in 200 μl S binding buffer [50mM HEPES pH 7.4, 150 Mm NaCl, 10 mM MgCl2, 0.05% NP-40] for 1 hr at 4 ℃ with gentle rotation followed by washing in S binding buffer (5 times for 10 min for each). Pre-blocked streptavidin beads were incubated with 6 to 10 μg of biotin-labeled RNA in 500 μl S binding buffer with 2 μl RNasin for 2 hr at 4 ℃. The RNA-bound streptavidin-agarose beads were added to the pre-cleared oocyte extract and incubated overnight at 4 ℃ with gentle inversion to allow protein binding. The RNP-bound streptavidin-agarose beads were washed 5 times with buffer (20 mM Tris-HCl pH 7.5, 0.25 mM EDTA, 6.25 mM MgCl_2_, 10% glycerol, 0.05% NP40), 10 min for each wash, and then incubated for 1 hr at 4 ℃ in 30 μl elution buffer (50 mM NH_4_HCO_3_, 8 mM biotin) with gentle pipetting up and down every 5 min to release biotin-RNP. All the steps involving collection of beads were done by centrifugation at 200 rpm for 1 min.

**Mass Spectrometry**

*Protein Preparation and Digestion*. Reagents and chemicals were obtained from Sigma Aldrich (St. Louis, MO) unless otherwise specified. Protein bound NA-beads were extracted, reduced, alkylated and digested with trypsin (*43*). Briefly, washed beads were resuspended in 6% SDS (sodium dodecyl sulfate) 100 mM TEAB (triethyl ammonium bicarbonate pH 8.5) 20 mM DTT (dithiothreitol) and heated at 95°C for 5 min. Then incubated at 60 °C for 15 min. Samples were alkylated with 2.5-fold molar excess of iodoacetamide for 20 min in the dark and quenched by addition of phosphoric acid to 1.2% final (v/v). A 7-fold volume excess of 90% methanol 100 mM TEAB was added to flocculate and each sample (including beads) was loaded onto a 200 µg-‘midi’ Strap (Protifi, NY,NY). The sample was washed twice with the same buffer and rehydrated with 2 µg sequencing grade trypsin (Promega, Madison WI) in 100mM TEAB and digested overnight at 37 ° C. Peptides were extracted by centrifugation, and desalted using an HLB 1 ml SPE (Waters, Beverly, MA) according to manufacturer’s instructions.

*LC-MS/MS*. LC-MS/MS was performed on a Q-Exactive HF (Thermo, San Jose, CA) running a TOP18 DDA. RAW files were searched and quantified using MaxLFQ and Andromeda within MaxQuant (v 1.6.2.3) against the current *Xenopus* FASTA sequence (Xenbase) concatenated with common contaminants (*44, 45*). False Discovery Rate was set to 0.01. Protein abundances were normalized against common abundant proteins co-associated with control RNA beads and data reduction was performed with a Benjamini-Hochberg corrected P-value to determine significantly enriched/reduced proteins (*46, 47*).

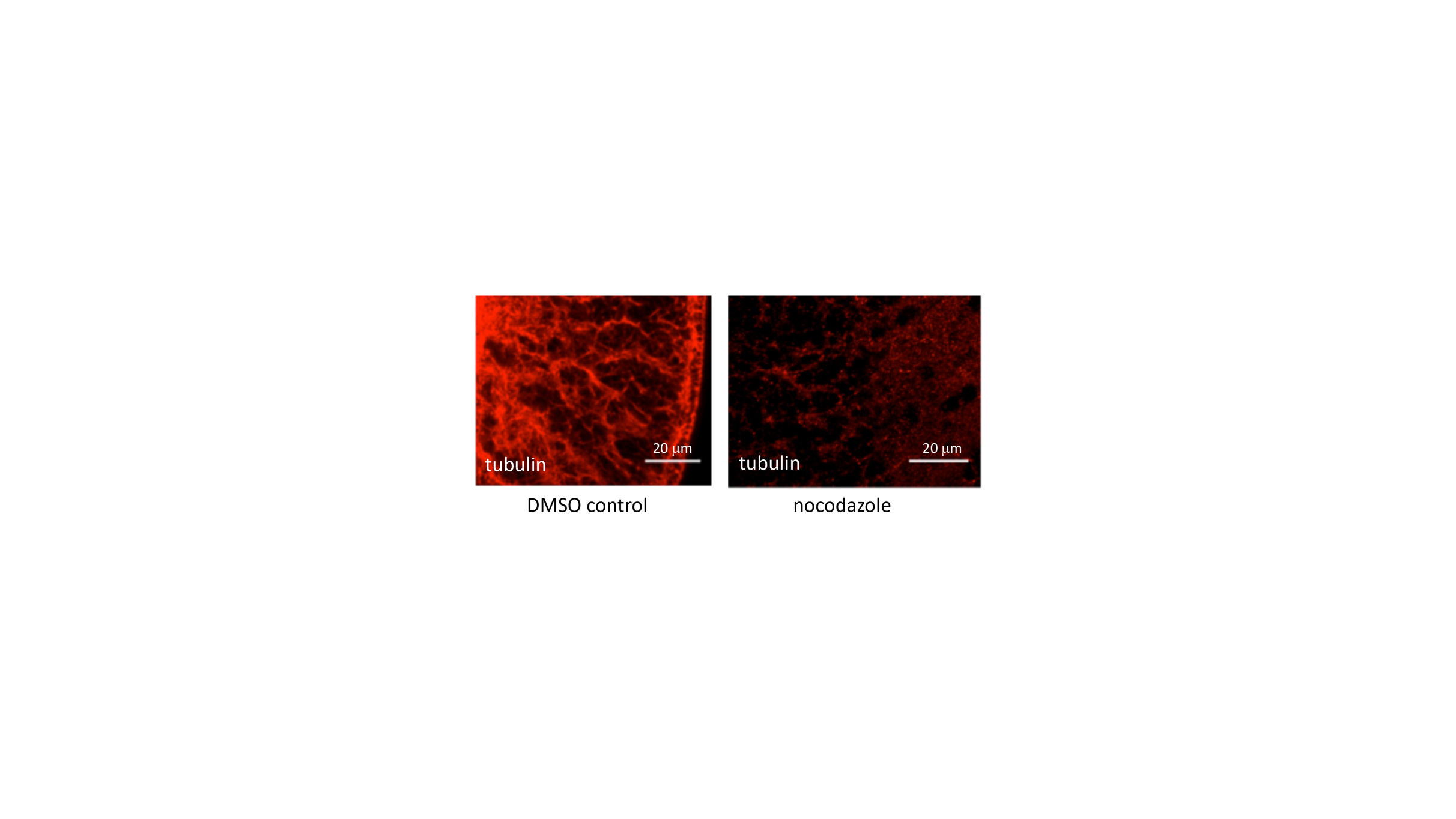

**Fig. S1.**

Depolymerization of microtubules with nocodazole. Oocytes were incubated with DMSO (vehicle control) or nocodazole (5 μg/ml) for 6 hr and then fixed for immunohistochemical staining with β-tubulin antibody.

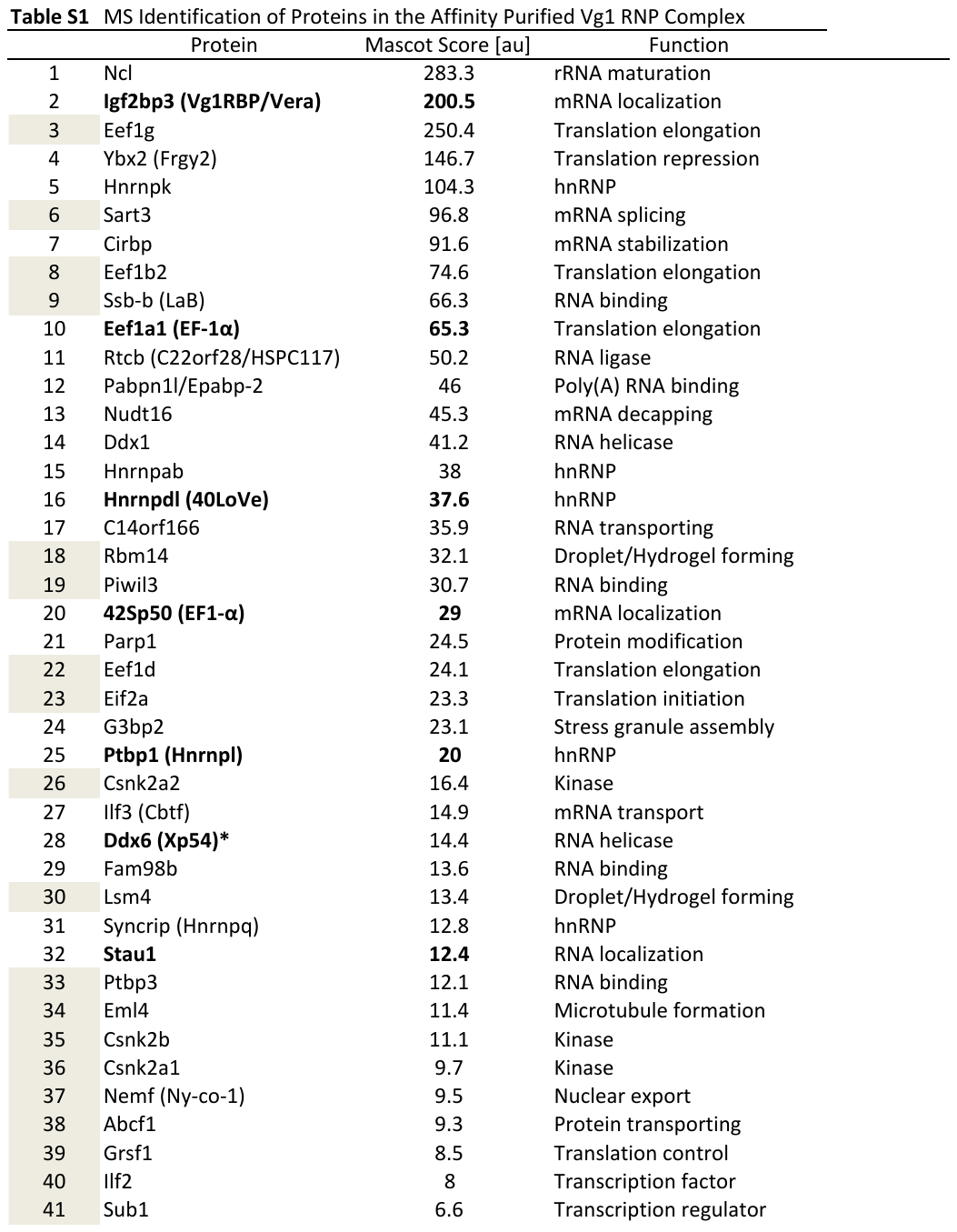

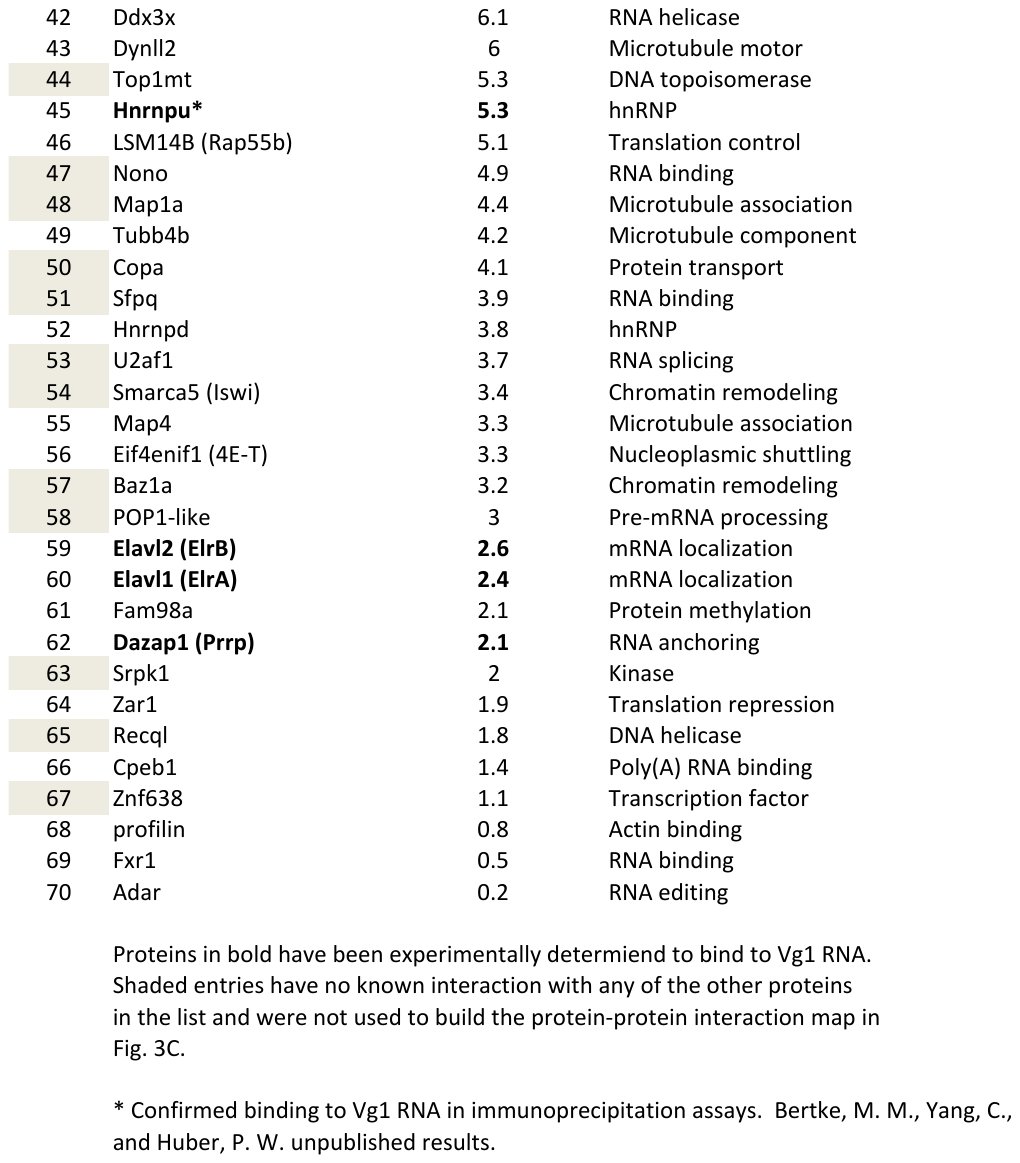
